# Interpreting phytoliths assemblages at chimpanzee (*Pan troglodytes verus*) nut-cracking sites in Bossou Forest, Guinea

**DOI:** 10.1101/2022.11.10.516074

**Authors:** C Phillips, K Almeida-Warren, MK Bamford

**Affiliations:** School of Anthropology and Museum Ethnography, University of Oxford; Evolutionary Studies Institute, University of the Witwatersrand

## Abstract

The nut-cracking behaviour of the chimpanzees of Bossou Forest has been long documented and studied in order to understand ultimate drivers for this form of durophagy by these apes. At sites in which they break open the nuts of the oil palm (*Elaeis guineaeensis*) on stone anvils with stone hammers, fragments of the tools as well as remnants of the nuts may be deposited into local sediments; however, they may become less visible at the macroscopic level as these sites are no longer used (become dormant). We build upon work that has been done to document this unique cultural heritage in West African chimpanzees by interpreting phytolith assemblages in sediments at active nut-cracking sites (used within the last two months). We compare these assemblages with those found in sediments of sites that have been dormant for ~10 years and sites where chimpanzees have not been observed to access and crack open oil palm nuts with stone tools. We predicted that larger assemblages of spheroid echinates, a phytolith associated with palms, would be found at active nut-cracking sites, however we found no statistical difference between total spheroid echinates (measured as total number found per gram of dry sediment) assemblages produced across active nut-cracking, dormant, or non-nut-cracking sites. This may have been due to small sample size (N≤6 sediment samples for each of the three sites) and so further sample collection and analyses are needed for inter-site comparisons. We also measured 2D area (μm^2^), perimeter and maximum diameter of spheroid echinates found in nut endocarps (shells) and leaflets from an oil palm frond. Intra-specific differences from all three measurements were found with these phytoliths being larger in the nut endocarp than the leaflet. This has implications for (re)interpreting the assemblages of spheroid echinate phytoliths at the three sites to determine if there is a greater productivity larger-sized spheroid echinates that fall within the size range of those measured for the nut endocarp (mean maximum diameter of 14.9 μm *versus* 7.8 μm for those found in oil palm leaflet). Finally, we argue for the importance of initialising and continuing the interpretation of phytolith assemblages in local sediments that are located near sites where directed plant input by non-human taxa has occurred, in this case, chimpanzees cracking open oil palm nuts using stone tools. This is important to refine our interpretation of phytolith assemblages where human and non-human taxa who use complex foraging strategies share ecological spaces.

## Introduction

### Durophagy in extant primates

The eating of nuts by our close primate relatives has received much attention to progress our understanding of their dietary ecology and material culture (Spagnoletti et al. 2012; McGraw et al. 2014), as well as to model aspects of diet, ecological niche, material culture, and cognitive abilities of human ancestors (Jolly 1985; Peters 1987; Yamakoshi 2008; Daegling et al. 2011; Bril et al. 2012). This form of durophagy has been observed in multiple primates including the Sooty mangabey (*Cercocebus atys*), bearded capuchin (*Sapajus libidinosus*), tufted capuchin (*Cebus apella*), Panamanian white-faced capuchin (*C. imitator*), western (*Pan troglodytes verus*) and Nigerian-Cameroon chimpanzee (*P. t. elliotii*), Japanese macaque (*Macaca fuscata*) and Burmese long-tailed macaque (*Macaca fascicularis aurea).* How they extract the kernal (endosperm) from their hard outer casing (endocarp) with tools (e.g., bearded capuchins using stone hammers and boulders as anvils: Langguth & Alonso 1997; Fragaszy et al. 2004; Panamanian white-faced capuchin using stone hammers and stone or wooden anvils: Barrett et al. 2018; and western chimpanzees using stone or wooden hammers and anvils: Sugiyama & Koman 1979; Boesch & Boesch 1983) or without tools (e.g., tufted capuchins hitting nuts directly on branches or against each other: Struthsaker & Leland 1977; Japanese macaques using teeth: Tamura 2020) continues to be documented. For primates who access and process nuts with tools, known as nut-cracking, this involves an individual selecting a nut (due to size, hardness and/or availability), and as mentioned, a stone/wooden hammer and a stone/wooden anvil (latter may be transportable or fixed e.g., tree root or branch, rock outcrops; Carvalho et al. 2008). The individual places the nut on the anvil and strikes the endocarp to access and feed on the endosperm. Successful individuals develop a greater spatial awareness between the hammer and nut and the nut and anvil (Matsuzawa 2001; Fragaszy et al. 2004). Indirect evidence e.g. fragmented refuse of nuts, indents or percussion marks on tools, foot and handprints near the nut remains or tools (Humle & Matsuzawa 2004; Morgan & Abwe 2006; Benitio-Calvo et al. 2015; Falótico et al. 2017), and direct observation of nut-cracking (Sugiyama & Koman 1979; Carvalho et al. 2008; Fragaszy et al. 2013; Luncz et al. 2017; Proffitt et al. 2018) have provided insight into how they access, process and ingests nuts. In turn, workers have addressed questions relating to social, cultural and ecological selection pressures that have driven primate durophagy and how an adaptation to feed on this hard-to-access food has shaped cognitive, anatomical and morphological traits (Carvalho et al. 2008; Wynn et al. 2011; Duarte et al. 2012; Fragaszy et al. 2013; Visalberghi & Fragaszy 2013; Sirianni et al. 2015; Fragaszy et al. 2017; Lombard et al. 2019). If we look at the Sooty mangabeys in TaÏ Forest, Ivory Coast as an example: they bite-through the endocarp of *Sacoglottis gabonensis* (Baill.) Urb. with their molars and premolars to access and eat the endosperm. These primates feed on this nut more frequently than any other food-item each month and across the year (McGraw & Daegling 2014) which has opened our questioning on how we categorise fallback foods (i.e., foods that are argued to be consumed when preferred foods are scarce) based on enamel thickness and tooth morphology (Daegling et al. 2011; McGraw et al. 2014). In addition, nuanced aspects of dietary related adaptations such as sex differences in ability to access harder nuts (Geissler et al. 2021), selection for gape ability and its impact on hard-food eating (Taylor et al. 2018) and the scavenging of plant matter, in this case on *Coula* (Baill.) and *Panda* (Pierre.) nut fragments left by chimpanzees and river hogs in TaÏ Forest (Bryndan et al. 2018) have also been explored. This example highlights the complexity in proximate and ultimate explanations of durophagy by primates (discussed further for chimpanzees below).

### Utilisation of the oil palm by primates (including humans)

The focus of our study is the nut-cracking sites created by chimpanzees in Bossou Forest, Guinea, (Sugiyama & Koman 1979, Matsuzawa 1994, Matsuzawa et al. 2001, Biro et al. 2003) but chimpanzee nut-cracking sites have been found across West Africa in Cameroon (Morgan & Abwe 2006), Sierra Leone (Whitesides 1985), and Liberia (Anderson et al. 1983; artefacts found for recent extirpated Ganta community: Smith et al. 2010); notably, nut-cracking by the North, South and East chimpanzee communities in Tai Forest, Ivory Coast has been observed across five decades (Struthsaker and Hunkeler 1971; Boesch & Boesch 1983; Luncz et al. 2018). As to how far reaching this foraging strategy originally occurred across the range of *P.t.* sp., and why this behaviour has only been documented in extant populations in West Africa remains unresolved. It has been suggested that natural barriers such as the N’Zo-Sassandra River in the Ivory Coast may have impacted the diffusion of this behaviour (Boesch *et al.* 1994; Humle and Matsuzawa 2004), or that it was innovated or devolved in pockets across populations (McGrew et al. 1997; Morgan & Abwe 2006). For context, dating of excavated nut-cracking sites with nut refuse and tool artefacts indicate durophagy and nut-cracking in western chimpanzees for at least 4300 years (Mercader et al. 2007 *vs*. 3000 years for bearded capuchins: Falótico et al. 2019); for humans, for at least 780,000 years (at Gesher Benot Ya’aqov, Israel: Goren-Inbar et al. 2002). As well as co-evolution of nut-cracking across the order, Primates, percussive technology origins in our lineage argued to predate the earliest stone tool artefacts dated to 3.3 million years (Harmond et al. 2015) allude that nut-cracking was a persistent foraging strategy across multiple hominin taxa (Wynn & McGrew 1989; Schroer 2012; Bril et al. 2015; Arroyo et al. 2016).

One nut that is accessed and processed by the Bossou chimpanzee community is that of the oil palm (*Elaeis guineensis* Jac q., family Arecaceae, subfamily Arecoideae; illustration of parts in Fig. 1). The oil from this nut is high in fatty acids and a good source of energy due to a high carbohydrate content (Atatsie et al. 2009; Dian et al. 2017). Oil palms are a single-branch monocot native to West and Central Africa but have since dispersed across parts of Africa (Robins 2021). Possible fossilised pollen of this palm occurrs in late Miocene sediments in Nigeria (Zeven 1964, 1972), as well as Late Holocene sediments and pottery artefacts across parts of West Africa (Sowunmi & Awosina 1991; D’Andrea et al. 2006; Garcin et al. 2018). Although likely to be a combination of factors, debate continues as to whether expansion of oil palm across Africa in the late Holocene was due to human activity (Hartley 1988), animal dispersal agents (Zons & Henderson 1989), that it is a pioneer species in disturbed areas, or in areas impacted by climate change such as the Dahomey Gap (Maley & Brenac 1998; Maley & Chepstow-Lusty 2001; Salzmann & Hoelzmann 2005; Blach-Overgaard et al. 2010). It is now cultivated worldwide across the humid tropics for oil which is extracted from the fibrous mesocarp of the fruit and the endosperm of the nut (Meijaard et al. 2018). Demand for palm oil has caused largescale deforestation and biodiversity loss (Fitzherbert et al. 2008; Koh and Wilcove 2008; Yapp et al. 2010; Vijay et al. 2016) with devastating impacts on the survivorship of primates (Wich et al. 2014; Linder & Palkovitz 2016; Supriatna et al. 2017), but like many palm species, faces the threat of decline with climate change (Murphy et al. 2021; Paterson 2021). Here, it should be noted that multiple primates feed on parts of the oil palm including the nut, flower, leaf and fruit (e.g., tufted capuchins: Struthsaker & Leland 1977; vervets (*Chlorocebus aethiops):* McGrew 1985; lesser white-nosed guenon (*Cercopithecus petaurista), Diana guenon (Cercopithecus diana):* Humle and Matsuzawa 2004; long-tailed macaque: Mathews et al. 2007; Panamanian white-faced capuchin: Estrada et al. 2012; Bornean orangutan (*Pongo pygmaeus):* Ancrenaz et al. 2014; Southern pig-tailed macaque (*Macaca nemestrina):* Ruppert et al. 2018, and chimpanzees (*P.t.* spp at various sites: Taï Forest Reserve, Ivory Coast: Boesch & Boesch 1983; Lopé Reserve, Gabon: Tutin et al. 1995; Ndoki Forest, Democratic Republic of Congo: Kuroda et al. 1996; Mahale and Gombe, Tanzania: Goodall 1986, Zamma et al. 2011). Chimpanzees and orangutans also build arboreal beds in oil palm trees (Sousa et al. 2008; Ancrenaz et al. 2014; Carvalho et al. 2015); whether due to necessity or opportunity remains unclear.

**Figure 1:**
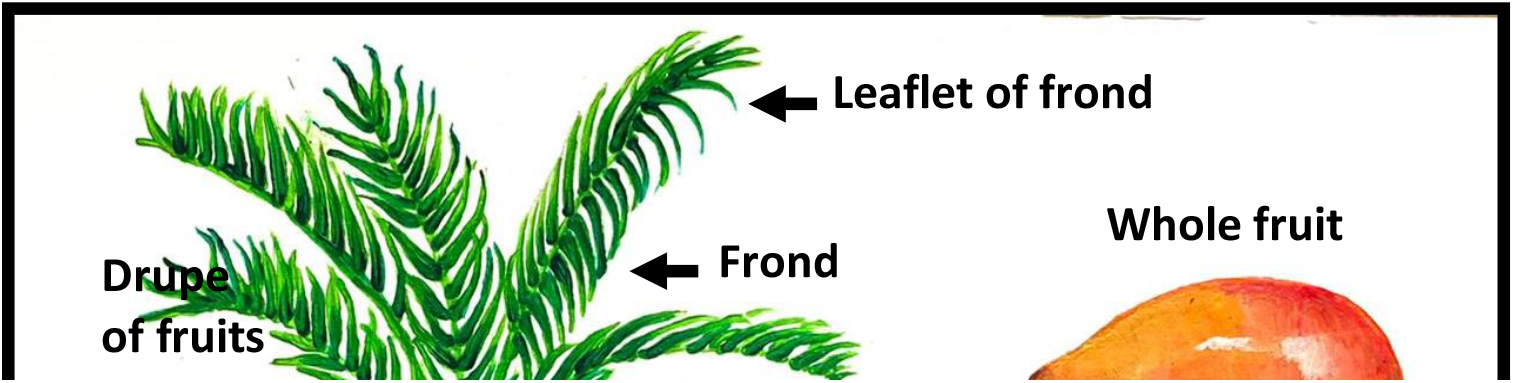
Parts of the oil palm (*Elaeis guineensis*). Illustration by Elodie Freymann.

**Figure 2:**
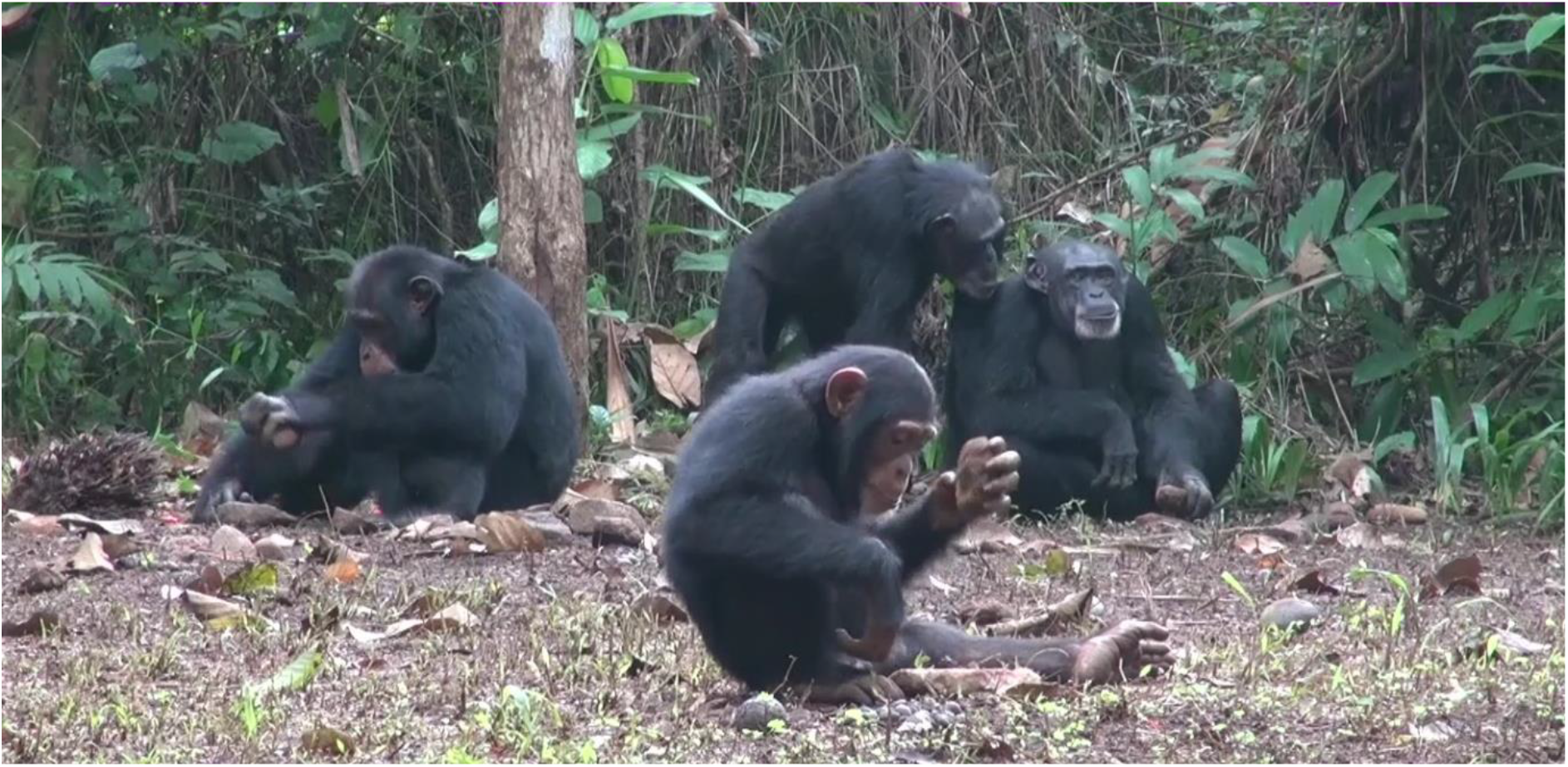
Bossou chimpanzees using stone tools (hammer and an anvil) to crack open nuts. Photo by Katarina Almeida-Warren.

The Bossou chimpanzee community not only feed on the oil palm nut, but also the leaflets and pith of the frond. Nuts and pith are eaten when preferred foods are scarce (Yamakoshi 1998) and the eating of these plant parts has become ever more prevalent in this ape community, partly driven by long-term anthropogenic activities (Hockings et al. 2009; Leblan & Soiret 2021). Many Bossou village inhabitants rely on swidden agriculture (i.e., cutting and burning of vegetation in a forested environment) (Yamakoshi & Leblan 2013) and the heterogeneric landscape that has been created has resulted succession of food resources in most habitats (Bryson-Morrison et al. 2016; Leblan & Soiret 2021), including oil palms that have thrived in disturbed local habitat. Many are heavily reliant on this resource (Yamakoshi 2011, 2013). Oil palms are also found in what has been referred to as ‘a buffer zone’ between core areas used by the chimpanzees and the villagers themselves, they have been considered ‘common property’ to both humans and chimpanzees, and historically, this inter-species resource sharing has facilitated co-existence between them to some degree (Yamakoshi 2005; 2011). The Bossou chimpanzees also access the youngest shoot tip (the apical meristem) located in the leaf bases of the crown shaft (Tomlinson et al. 2011). The chimpanzees first laboriously pull up older shoots and feed on the petiole and leaf base of these. This creates a hole in what looks like a vertical cylinder, and they access the flesh of the newest tip in the crown shaft by pounding it with one of the pulled-out, older shoots (Yamakoshi 2011). This behaviour is known as pestle-pounding, so far, only observed for this ape community and a neighbouring community in Kpala, Liberia (Sugiyama 1994; Yamakoshi & Sugiyama 1995; Yamakoshi 2011; Ohashi 2015). As well as the Bossou chimpanzees, neighbouring chimpanzee communities in Diéké and Yealé, (Humle and Matsuzawa 2004; Matsuzawa et al. 2008; Humle 2011; Carvalho 2013) also practice nut-cracking of the oil palm nut; however, not all seem to (Humle & Matsuzawa 2001; Koops et al. 2013; Phillips C personal observation for Nimba community).

Nut-cracking behaviour by the Bossou chimpanzees has been observed for over four decades (Sugiyama & Koman 1979; Biro et al. 2010; Bryson-Morrison et al. 2020) and has contributed to revealing complexities in understanding proximate and ultimate explanations for the practice of durophagy of nuts by these apes. The châines opératoires of this phenomenon (Carvalho et al. 2008), as well as ontogeny and proficiency of nut-cracking (Biro et al. 2006), transportation and re-use of tools (Sakura & Matsuzawa 1991; Carvalho et al. 2009), tool selection and function, forms of locomotion used and laterality of nut-crackers (Humle & Matsuzawa 2009; Carvalho et al. 2012) have all been studied to understand proximate mechanisms to consider ultimate explanations for ecological, biological adaptations e.g. nutrition (Bryson-Morrison et al. 2020), mental mapping of availability of nuts and tools as observed for Taï Forest communities (Boesch & Boesch 1984), as well as cultural and social selection pressures (Bossou Forest: Biro et al. 2003; Matsuzawa et al. 2008; Taï Forest: Boesch et al. 1994: Luncz et al. 2012) for increased fitness and reproductive success. A key observation by Mercader et al. (2002) from their excavations of chimpanzee nut-cracking sites in TaÏ Forest was that *“chimpanzees left behind stone and plant refuse_that accumulated in specific loci”.* Our work aims to build on this observation by exploring further the plant refuse accumulated from nut-cracking by the Bossou chimpanzees; however, at the microscopic level rather than the macroscopic level which may reveal greater insight from localised, decomposed plant refuse no longer visible to the naked eye. We achieved this by analysing oil palm phytoliths produced in specific loci sediments at chimpanzee nut-cracking sites in Bossou Forest (further details below).

### Why phytoliths? Overview and application of phytoliths across Africa

Phytoliths are opal silica microfossils (‘phyto’ meaning plant and ‘lith’ meaning stone: Baker, 1959) and are formed when aqueous monosilic acid (Si(OH)4) enters the plant during water, nutrient and other mineral uptake and is then deposited within cell lumina, in the cells’ wall or in between cells. The deposited silica then solidifies producing silica plant-cell-structure formations to varying degrees of silicification (Madella & Lancelotti 2012) during evapotranspiration. Phytoliths encountered in soils have normally accumulated due to deposition by local, phytolith-producing plants as they decompose, although some translocation can occur vertically and horizontally by water and air movement, and translocation distance can vary between different phytoliths (Fishkis et al. 2009, 2010; Lui et al. 2019). Organic carbon can be occluded (i.e., blocked) in phytoliths which can impact soil carbon sequestration (Par & Sullivan 2005) and deposited phytoliths can breakdown or dissolve in soils due to many factors including pH, surface bulk ratio (i.e., surface area to volume ratio of phytolith shapes; Cabanes et al. 2011; Cabanes & Shahack-Gross 2015; Strömberg et al. 2018). Such factors can help maintain equilibrium in concentrations of silica in soils (Farmer et al. 2005), but they can also evade these processes and therefore, remain preserved and undisturbed in soils and paleosols worldwide (Runge & Fimbel 2001; Piperno 2006; Cabanes et al. 2011). Furthermore, phytoliths can survive charring, mineralisation, oxidisation (unlike pollen in more tropical regions) and when ingested, mastication, digestion and defaecation (Cabanes et al. 2011; Hart 2016; Cabanes 2020). This, along with the fact that they can be distinctive in chemistry, size and morphology (known as morphotypes: Rovner 1971; Piperno 1988; Piperno 2006; Hodson 2016) allow workers to diagnose and associate phytoliths to plant type (e.g., grass), to family and genus, and with more recent morphometric work, potentially to species (Hart 2016; Cai & Ge 2017; Hošková et al. 2021). Additionally, phytolith indices have been created using combinations of phytolith morphotypes to indicate aridity and climate (Twiss et al. 1992), and tree cover (Alexandre et al. 1997; Barboni et al. 2007). Thus, phytoliths are a powerful palaeoecological tool to reconstruct diet and environments of extant and extinct taxa (Aleman et al. 2012; de Silva Neto et al. 2020) including humans and our past activity (Pearsall 1994; Piperno 1994; Jiang, 1995; Runge 2001; Madella et al. 2002; Wanget et al. 2003; Barboni et al. 2010).

Phytoliths have been studied across Africa, but relatively more recently compared to work done on other continents (Piperno 2006). A key focus has been to document phytolith morphotypes across African flora and create reference libraries of multiple morphotypes that are associated with or indicate vegetation structures and habitats to assist reconstruction of modern and past environments across Africa (Runge & Runge 1997; Runge 1999; Mercader et al. 2000; Bremond et al. 2008; Barboni & Bremond 2009; Aleman et al. 2012; Bremond et al. 2017; Bremond et al. 2020). For palm phytoliths, work has focused on morphotypes in *Hyphaene* and *Pheonix* at African archaeological and modern sites, particularly on their presence, preservation, and abundance in sediments due to taphonomic processes and size of phytoliths (Olduvai Gorge, Tanzania: Bamford et al. 2006; Albert et al. 2009, Barboni et al. 2010, 2014) as well as their presence in relation to surface groundwater levels (multiple sites in East Africa: Albert et al. 2015; Barboni et al. 2019). This work builds upon phytolith analyses of multiple palm species globally (e.g. phytolith variability in Amazonian palms and their use by humans: Piperno 1989; Morcote-Rios et al. 2013, 2016; Witteveen et al. 2022; morphotype analyses of palms at archaeological sites in Papua New Guinea and Rapa Nui: Delhon & Orliac 2010; Fenwick et al. 2011; Bowdery 2015). Such attention to palms is unsurprising as they are high phytolith producers (Hodson et al. 2006, 2015), their phytoliths are relatively robust to chemical and physical processes pre- and post-deposition due to factors such as having a surface bulk ratio >1 (Barboni et al. 2019). Furthermore, they have relatively distinct taxonomic phytolith morphotypes, including the spheroid echinate. As described and identified using Neumann et al.’s (2019) Code for Phytolith Nomenclature (IPCN) 2.0, these phytoliths have a basic shape representing a sphere, and the echinate denotes the surface texture and ornamentation of having conical projections that “vary in size which can be pointed or rounded” (Fig. 5). For spheroid echinates associated with the palm family, Arecaceae, they range in size of 6-25 μm (Piperno 2006; Benvenuto et al. 2015), and their conical formations are generally large and closely distributed (Neumann et al. 2019).

As well as building upon Mercader et al.’s (2000) observation at Taï Forest to consider the microscopic level of oil palm nut refuse, we also highlight a component of plant input that requires further attention; that is *directed* plant input that results from complex foraging behaviours of non-human taxa, such as accessing, processing of foods with lithic and perishable tools, which we argue is not equivalent to natural or anthropic plant input (Zurro et al. 2016). Directed plant input caused in this case, by the chimpanzees repeatedly crushing nut endocarp and possibly the kernel (if the nut is hit too hard) with a stone tool could enter sediments and as Madella and Lancelotti (2012) argue for anthropic plant input could, *“produce phytolith assemblages that are normally much larger than those produced from the natural vegetation* [i.e., natural input] *or the phytolith soil bank”.* As remarked by these workers, we acknowledge that there is a spectrum of anthropic plant input that depends on specific practices across human societies. This would, therefore, have implications for phytolith assemblages produced, for example, cracking open nuts of oil palms might produce a ‘weaker’ (i.e. produce phytolith assemblages that are normally less large) anthropic plant input than heating up of oil palm over an open fire (with palm parts as fuel), but if the former was repeated at a highly localised sites, this deserves consideration when interpretating phytolith assemblage in associated sediments. Work on the impact of bioturbation by fossorial insects on phytolith production in sediments indicate the importance to understand how multiple non-human taxa influence sediment phytolith assemblages (Hart 2003; Jouquet et al. 2020). For primates, such work would acknowledge the mutual roles that they play in shared ecological spaces with humans (Fuentes 2012). Our aim was to analyse oil palm phytoliths at: 1) active sites where nut-cracking has been observed in the last five years; 2) dormant sites where nut-cracking has not been observed for 10 years); and 3) non-nut-cracking sites where this form of durophagy has never been observed to determine if resultant accumulation of oil palm nut refuse from nut-cracking by chimpanzees provides a phytolith signature in “specific loci” (Mercader et al. 2000). We tested H1: Production of spheroid echinate phytoliths will be larger in localised sediments collected in close proximity to visible remains of cracked oil palm nuts (within 10cm of remains). We predicted a larger production of spheroid echinate phytoliths at the active nut-cracking sites based on the assumption that oil palm nut refuse fragments pre- and post-deposition, would be present in greater frequency in localised sediments.

We also endeavour to build upon work undertaken to reconstruct diets of extant primates using micro remains such as phytoliths (reviewed Henry 2012). Diet composition estimates using phytolith reference libraries from plants eaten by extant primates been recently explored. Phytolith assemblages have been correlated with primate enamel thickness (Power et al. 2015), and interpretations of phytolith assemblages in faeces of east and west African chimpanzees (Phillips & Lancelotti 2014; Power et al. 2021) have revealed components of diet that are less detectable at the macroscopic level. Relevant for this study and for future work is Power et al.’s (2021) findings for the nut-cracking chimpanzees of Taï Forest; oil palm phytoliths were “overrepresented” in dietary composition estimates based on analyses of their faeces and dental calculus. These apes access endosperms of oil palm nuts as well as non-palm nuts of *Coula* and *Panda* through nut-cracking; it was suggested the spines of the spheroid echinate phytoliths from oil palms could perhaps embed themselves more easily in dental plaque versus phytoliths with a smoother (psilate) surface (Power et al. 2021). Such work is vital for continued efforts to create high-resolution dietary comparatives of non-human taxa using phytolith analyses, yet ethnographic comparatives beyond consumption remain lacking – in this case, directed oil palm nut input into sediments associated with nut-cracking tool-use behaviour of western chimpanzees.

Finally, we aim to complement work done on the behavioural and technological landscape of the Bossou chimpanzees which provides support for the favoured places hypothesis for patterns of landscape use in relation to nut-cracking (Almeida-Warren et al. 2021) and contribute to the continued documentation of the behavioural ecology and unique cultural heritage of this ape community. Few individuals remain (see methods below) and their sharing of ecological space with the people of Bossou is also motive to document and interpret phytolith assemblages from directed plant input of these non-human primates. Anthropic plant input could potentially result in directed input by these primates being overlooked.

## Methods

### Study site

Situated near the border of Ivory Coast and Liberia, Bossou Forest surrounds the village which is located in the sub-prefecture of Lola, South-Eastern Republic of Guinea (7°38’71.7’N - 8°29’38.9’W) at an attitude of 550m above sea level (Humle 2011). In terms of geology, in this southern part of the West African Craton, the Archaean rocks (dated 2.8Ga) are defined by Tonalite-Trondhjemite-Granodiorite (TTG) gneisses (Rollinson 2016). There are also supracrustal belts containing greenstone, and iron formation sequences dated 2.7Ga. Paleoproterozoic (dated 2.3Ga) reworking of the Archaean rocks are also found in the form of an intrusion of Macenta Batholith (White & Leo 1969), which consists of coarse-grained, igneous granitoids (Rollinson 2016). Leptosols which are typically shallow soils over hard bedrock are recorded for this area, but a detailed analyses of soil type is yet to be done for Bossou and possibly there could be anthrosols i.e., soils that have been heavily modified from human activity (WRB 2022). Analyses of a section of soil in Bossou indicated moderately acidic soil (pH range: 4.9-5.7) with sand to silt-sized particles (SM Table 1). Application of the feel method on-site, indicated a loam soil texture (Brady & Weil 1996). Furthermore, the yellow-red hue (5-7.5YR Munsell Colour Chart) of the soil reflects iron oxidation. Soil temperatures are isomegathermic (Jones et al. 2013) which means ≤5°C difference between the dry season (November to February), and the wet season (March to October). In terms of topography, the home range of the Bossou chimpanzees includes four hills (Gban, Gbouton and Guein) with elevations between 70-150m (Fig. 3). Vegetation across their range is a mosaic of primary (found only on Gban), secondary and gallery forest across 6km^2^. Secondary forest at different successional stages is the most prevalent habitat (Bryson-Morrison et al. 2016) where patches of fallow, coffee plantations and cultivated areas of varying degrees are found (Sugiyama & Koman 1992; Hockings et al. 2009). Dominant trees include: *Albizia zygia* (DC.) J.F.Macbr., *Sterculia tragacantha* Lindl. and various species of *Ficus* spp. *Thaumatococcus daniellii* (Benn.) Benth. and *Aframomum latifolium* K. Schum. are terrestrial herbaceous vegetation found across their home range and are eaten by the Bossou chimpanzees (Bryson-Morrison et al. 2016). Oil palms have been recorded on three of the hills of the forest. Palms located on the top of the hills are not used or cut down by humans due to traditional beliefs (see below; Yamakoshi & Leblan 2013); Schnell (1946) suggests that the Bossou chimpanzees have been key dispersal agents for these particular oil palms.

**Figure 3:**
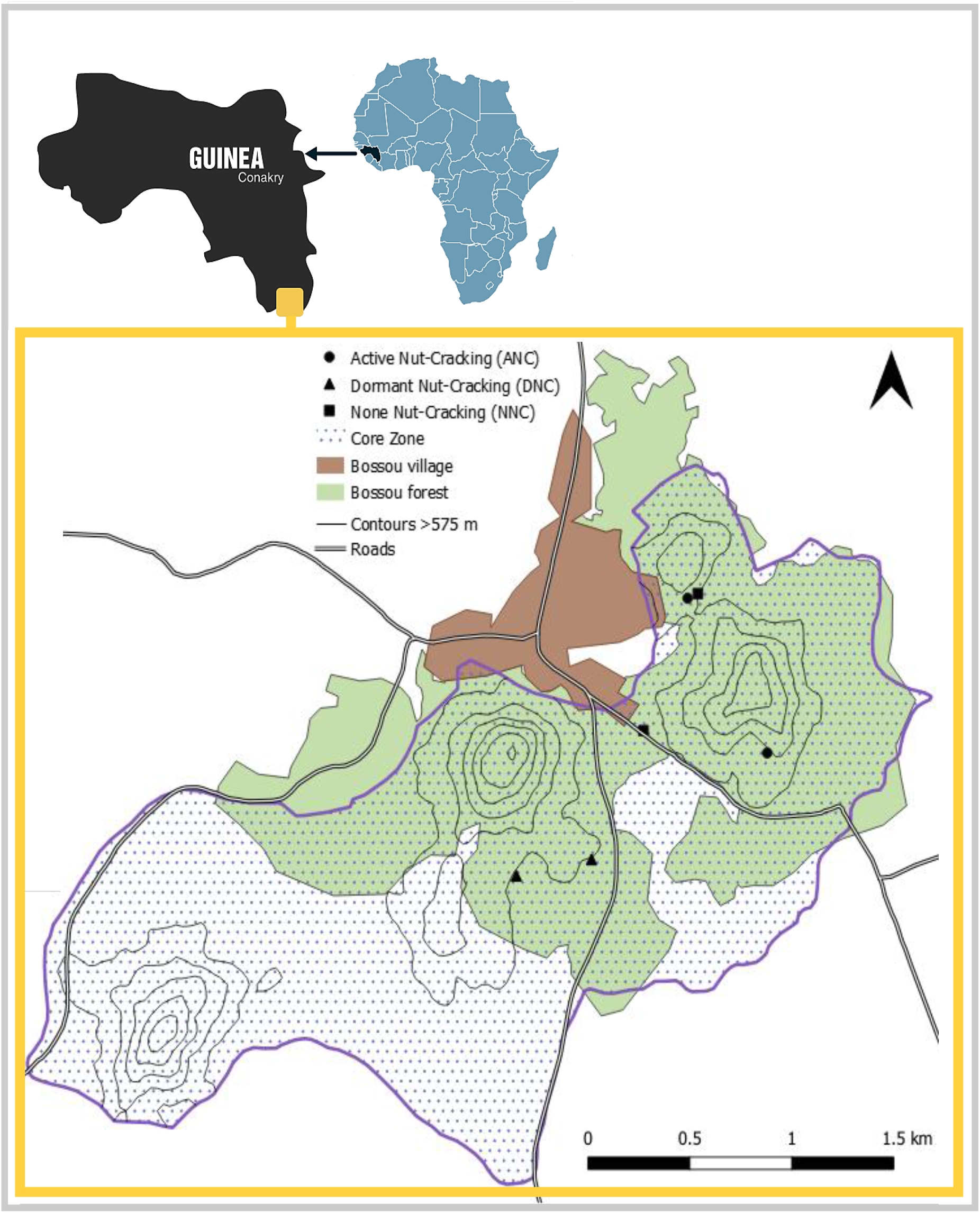
Map of Bossou Forest in Guinea with location of active nut-cracking sites (ANC) which had been used within 2 months of sampling, dormant nut-cracking sites (DNC) which had not been observed to be used at a nut-cracking site by the Bossou chimpanzees for ~10 years and non-nut-cracking sites (NNC) which have not been observed to be used as a nut-cracking site. Map also depicts home range of the Bossou chimpanzees (purple) which was adapted from Humle (2011). Contours of hills (thin black lines), village (brown), forested area which includes primary, secondary and gallery forest, as well as some cultivated areas (green) and roads adapted from ©OpenStreetMap and NASA (2013). Map of Africa and Guinea created on Visme.

### People and chimpanzees

The Bossou chimpanzees have been studied for almost 45 years with longer-term studies beginning in 1976 (Sugiyama and Koman 1979; Sugiyama & Fujita 2011). During this time, community size has been no more than 23 individuals, but they have experienced significant losses due to disease since 2003 (Humle 2011: Matsuzawa 2020) and current numbers are less than 10. They are fully habituated and so tolerate select human observers at close proximity (Williamson et al. 2011), allowing diurnal follows to be undertaken within their 15km^2^ home range. These apes have a long tradition of sharing ecological and social space with the people of Bossou. Currently, the village comprises of people of multiple backgrounds and ethnicities including Muslim Fula, Malinke, Sus, Yakuba, Kono and Manon (Granier & Martinez 2011; Yamakoshi & Leblan 2013). The traditional animist religion is practised by some and the Bossou chimpanzees have not been hunted for meat due to a long-standing ban initiated by clans as well as the belief that the chimpanzees are ancestral inhabitants who have changed form (Yamakoshi 2006; Yamakoshi & Leblan 2013). Chimpanzees mostly utilise and inhabit the upper forested parts of three of the hills in their home range, however, lower down swidden agriculture is practiced, and resource competition is observed through crop damage by the chimpanzees, including oil palm fruits (Hockings et al. 2009). Whether sustainable co-existence between the Bossou people and the last remaining chimpanzees is achievable remains to be seen. This is ever more imperative when savannah and further patches of plantations and fallow surround Bossou Forest and the village where *Pennisetum purpureum* Schumach., *Harungana madagascariensis* Lam. ex Poir., and *Sarcocephalus latifolius* (Sm.) E.A.Bruce are dominant (Morimura et al. 2014). Biogeographically, the nearest continuous primary and secondary montane forest, gallery forest and neighbouring chimpanzee communities are southeast of Bossou and are 6km away in the Guinean Nimba Mountains which is part of a Strict Nature Reserve and UNESCO World Heritage Site (Humle 2011).

### Taphonomy of phytoliths

Despite their robusticity, oil palm phytoliths with varying degrees of silicification may have been deposited into sediments through natural input (e.g., parts falling of the palm, parts dropped when animals are feeding oil palm or when the palm dies and decomposes). With directed input of oil palm phytoliths from nut-cracking activity of the chimpanzees, the percussive action to open the nuts with stone tools could have caused damage to some of the phytoliths pre-deposition. Regarding post-deposition, recent work indicates that rate of dissolution of phytoliths in oil palm fronds is lower than in phytoliths detected in rainforest litter (von der Lühe et al. 2022) which may, albeit tentatively, provide support for greater survivorship of oil palm phytoliths to occur in sediments of the primary, secondary and gallery forests of Bossou. As mentioned, particle size appears to indicate sand and silt-sized particles in the shallow soils of this forest which might have implications for the movement of phytoliths of ≤62-2 μm moving downwards to deeper strata depth in sediments. The impact of hitting nuts on stone anvils with stone hammers on such phytolith movement in surrounding sediments remains unknown but warrants further investigation if spheroid echinates of palms could fall between these particles based on their estimated size of 6-25 μm. Bioturbation by fossorial insects such as *Macrotermes* ssp. and *Dorylus* spp. are also a consideration for interpretation of phytolith assemblages in Bossou Forest; for this study, we did not measure how close nests of both were to the palms and sediments sampled. Both insects are eaten by the Bossou chimpanzees using perishable tools (Humle 1999; Humle et al. 2009) and their locality to sampled areas in future warrants attention.

### Sample collection

In 2018 KAW collected sediment samples for phytolith analyses from six palms within the Bossou chimpanzee home range (Fig 3): two were palms where chimpanzees actively nut-cracked in the last two months (ANC), two dormant palms, where they had not been observed nut cracking since 2009 (DNC) and two where the chimpanzees had never been observed to nut crack as our control samples (NNC). Previous studies have used a 2m (Koop et al. 2013; Almeida-Warren et al. 2021), or 5m radius at nut-cracking sites (Humle & Matzusawa 2004); we applied the shortest radius as this is where we encountered tools and oil palm nut remains close to tools at active nut-cracking sites (Fig 4 a-b). Maximum sediment sample depth was determined using a Van Walt Auger (57mm diameter) which indicated a maximum soil depth of 23cm before the auger hit bedrock; hence, KAW collected at depths of 0-5cm, 5-10cm and 10-15cm by creating a small soil profile and removing samples using deposable spatulas (N=17: Fig 4c). Two samples of oil palm nut endocarp and two leaflets from a frond were collected by CP in 2013. All samples were placed in double, pre-labelled plastic zip-lock bags. This work was permitted under the Direction Nationale de la Recherche Scientifique et Technique and the Institut de Recherche Environnementale de Bossou, Republic of Guinea. Sample size of plant and sediments collected was minimised to reduce disturbance to the Bossou chimpanzees.

**Figure 4:**
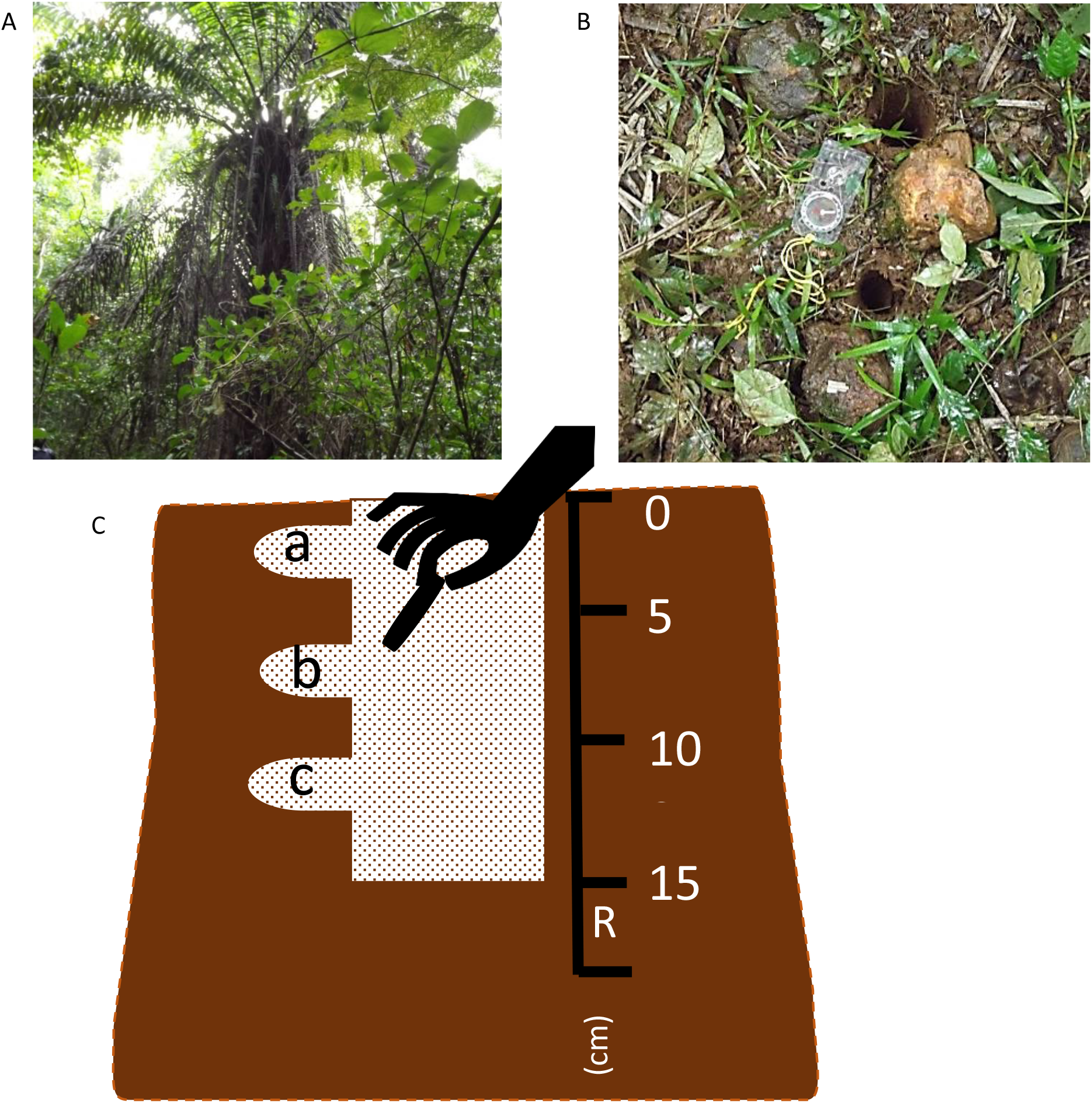
One of the oil palm trees (a) in which sediment samples were collected and (b) soil sample collection sites located in the vicinity of a stone tool and oil palm nut remains. Procedure for soil sample collection at different depths (c).

### Sample processing

We separated collected plant parts and prepared and extracted phytoliths from these four ashed plant samples (550°C to burn away organic matter in a muffle furnace) using Lancelotti’s (2010) dry ash extraction method (protocol outlined Phillips & Lancelotti 2014). For sediment samples, we applied Albert et al.’s (1999) extraction method but adapted it by: 1) running samples at 450°C in a muffle furnace rather than oven drying to remove organic matter; 2) after heating samples in boiling water to remove phosphates and calcium carbonates, using Jackson’s (1956) equation for clay fractionation in the centrifuge which resulted in running samples at 1000rpm x 4 mins x 30 cycles; 3) using sodium polystungstate solution of 2.3g/ml density to extract phytoliths; 4) running at 3000rpm for 10 mins instead of 5 mins until there was no more visible mineral particles in the supernatant. We then applied their AIF/g equation to provide a standardised total weight of acid insoluble fraction (containing extracted spheroid echinate phytoliths) per gram of dry sediment and dry plant for each sample (SM 2). Mounting medium used for slides and as a sealant was Entellan™. Phytolith processing of the four plant samples was done in the Palynology and Phytolith Laboratory at the Evolutionary Studies Institute, University of the Witwatersrand. The remaining samples were processed in the Oxford Long-term Ecology Laboratory at Dept. of Biology, University of Oxford.

### Phytolith taxonomy and analyses

We observed extracted phytoliths from both plant and sediment samples under a transmitted light MI5B-P Swift microscope at 40x and any photographs were taken using D-Moticam 10 camera and on Motic Images Plus 2.0. The description and identification outlined in Neumann et al.’s (2019) International Code for Phytolith Nomenclature (IPCN) 2.0 was used for this study of spheroid echinate phytoliths associated with oil palm. We counted complete spheroid echinate phytoliths along nine random vertical transects per slide, resulting in a total of 351 view fields (39 view fields per transect) based on a minimum count of 350 for valid sample material representation per slide (Van der Veen & Fieller 1982). This included the individual counting of these phytoliths within parts of plant cells where they were still articulated. These phytoliths are normally associated with the stegmata which is a cell found near vascular bundles and fibres in various parts of the palm (Schmitt et al. 1995; Neumann et al. 2019).

During the counting of the spheroid echinate phytoliths in both the nut and leaflet samples, there appeared to be a visible size-difference for each palm part. We therefore took an approximate measure of the 2D area (μm^2^) (Hart 2016), perimeter (μm) and approximate maximum diameter (μm) (Benvenuto et al. 2015) of 30 spheroid echinate phytoliths from a nut and leaflet sample to determine if this could give a preliminary indication to intra-species size differences for this morphotype in the oil palms at Bossou Forest (SM 3). We used the ‘irregular’ function on the Motic Images Plus 2.0 to draw around the echinoid surface of each phytolith to determine an approximated 2D area and perimeter, and the ‘line’ function for approximated maximum diameter of measured phytoliths (Fig. 5).

**Figure 5:**
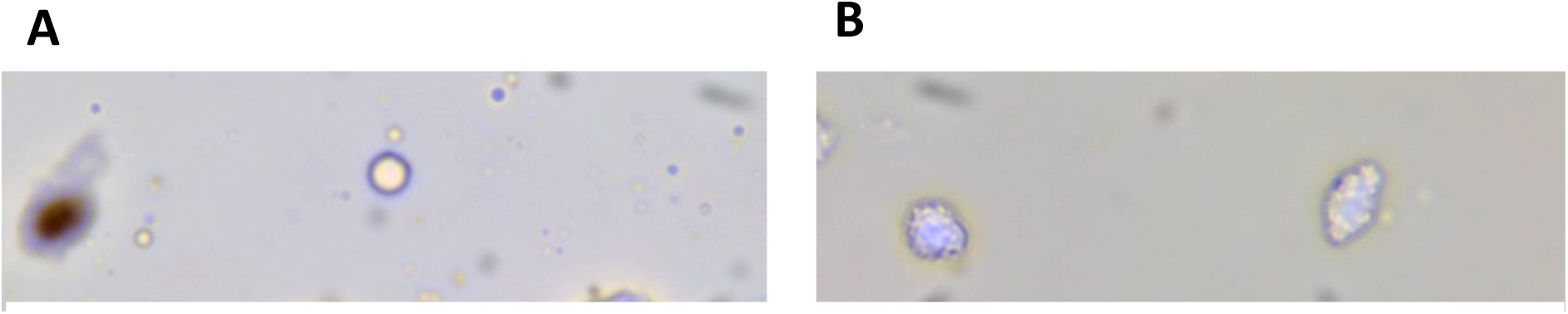
Spheroid echinates detected in (a) nut endocarp and (b) leaflet of the oil palm (*Elaeis guinaeensis*). Approximated measurement of 2D area (μm^2^), perimeter (μm) and maximum diameter of phytolith (μm) taken on Motic Images Plus 2.0.

### Data Analyses

To test if there were a larger production of spheroid echinate phytoliths in localised sediments at active nut-cracking sites (ANC) versus non-nut-cracking (NNC) and dormant nut-cracking sites (DNC), we carried out a Kruskall Wallis Test to compare mean AIF/g of collective sediments at ANC sites, DO sites and NNC sites as one-way ANOVA normality assumption was not met. To compare size variation between nut and leaflet spheroid echinates, a Wilcoxon signed rank test was performed for 2D area (μm^2^) and for the perimeter (μm) and maximum diameter (μm), a paired t-test (two-tailed). Analyses was done on (STATA IC 15.0), Shapiro-Wilk tests were done for normality due to sample size (<50) and α = 0.05 (SM 3).

## Results

Total spheroid echinates per gram of dry sediment (AIF/g) in ANC, DNC and NNC sediment samples are outlined in Figure 6. From this, it would appear that the AIF/g of spheroid echinate phytoliths across the different depths at the two non-nut-cracking sites appear relatively uniform, but no statistical analyses could be done on possible intra- and inter-site differences with this small sample size. No statistical difference was found between ANC, DNC or NNC sites for the combining of sediment sample AIF/g values at each site (*X*^2^ = 0.982, df = 2, p = 0621). Total spheroid echinates in AIF/g nut endocarp was 4876 and in leaflet it was 53529; many of the spheroid echinates within the leaflet remained articulated in silica skeleton compared to those detected in the nut endocarp. This aligns with other palm phytolith work on intra-specific productivity, with leaflets having greater numbers of palm phytoliths (Delhon & Orliac 2010).

**Figure 6:**
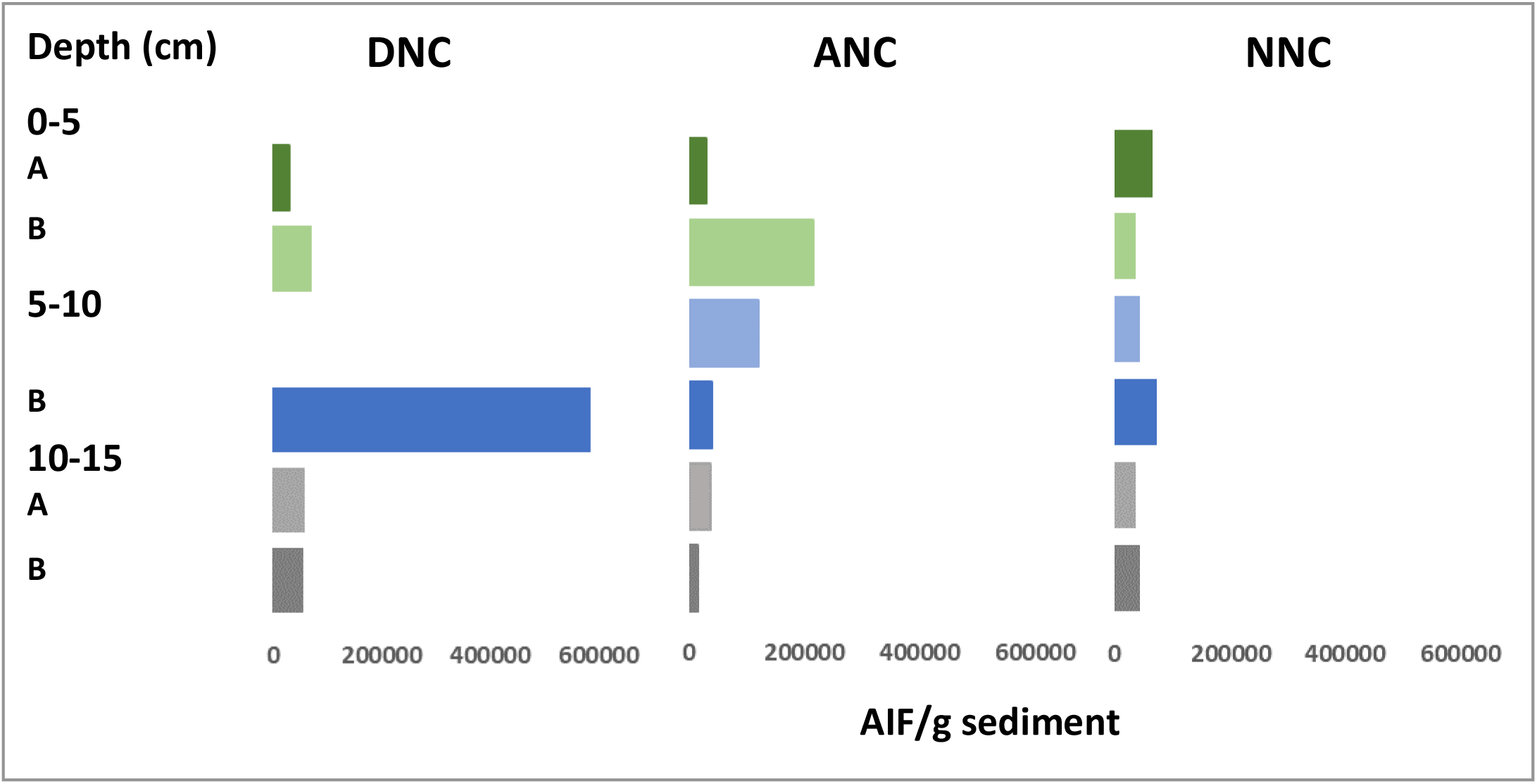
Bar graphs indicating total spheroid echinates recorded per gram of dry sediment (AIF/g) at oil palms (A + B) for each of the type of site: dormant nut-cracking site (DNC), active nut-cracking (ANC) and non-nut-cracking site (NNC). The green bars indicate AIF/g at sediment depth of 0-5cm, blue bars for 5-10cm and grey bars for sediment depth 10-15cm.

For 2D area of nut and leaf spheroid echinates ranged between 18.0-404.6 μm^2^ and 17.9-83.3 μm^2^ respectively and were statistically different (Wilcoxon signed rank test: z=4,960, p <0.001, N=30, effect size 0.85). Their perimeter ranges were 20.7-80.7 μm and 18.0-38.7 μm respectively and again were statistically different (Paired t-test (two-sided): T=10.79, df = 30, p <0.001, effect size 1.97, power 0.754). Maximum diameter of spheroid echinate phytoliths between the plant parts were also statistically significant (Paired t-test (two-sided): (Paired t-test (two-sided): T=9.353, df = 29, p <0.001, effect size 1.71, power 0.754); maximum diameter range: 5.0-25.1 μm for nut and 4.9-11.6 μm for leaf. Generally, these approximate maximum diameter ranges align with the 6-25 μm for spheroid echinates of Arecaceae proposed in Neumann et al. (2019), on two, especially with mean maximum diameters of 14.9 μm and 7.8 μm for the nut and leaflet sample respectively.

## Discussion

This study has provided initial findings to interpret phytolith assemblages from directed plant input by non-human taxa, in this instance as a result of the Bossou chimpanzees cracking open oil palm nuts using stone tools. Our prediction that there would be a larger production of spheroid echinate phytoliths in the localised sediments in close proximity to stone tools and oil palm nut remains at active nut-cracking sites was not met. It could be that the directed plant input by these apes may produce a relatively small phytolith assemblage as argued by Madella and Lancelotti (2012) for hunter-gather *versus* agrarian societies in relation to interpreting phytolith assemblages from anthropic plant input. However, our sample sizes were small (N=5 combined sediment samples for DNC; N=6 for combined sediment samples NNC; and N=6 for ANC sites). This has implications for effect size and type 1 error (Makin & Orban de Xivry 2019), and so a next step will be to increase sample size. First, in terms of the number of oil palms that localised sediment samples are collected from and second, to explore the phytolith productivity in sediments of the various depths, which although were collected in this study, were not fully analysed due to limitation of sample size.

Almeida-Warren et al. (2021) highlight that there are multiple oil palm trees (~40) that have been utlised by the Bossou chimpanzees to access and process oil palm nuts, and so there is great potential to explore and further interpret phytolith assemblages produced from nut-cracking activity by these apes. Further sampling of sediments at oil palms not used by the chimpanzees for nut-cracking are needed as continued control samples, but this will require some consideration in terms of their location and historical use. Oil palms in this region are under various stages of succession due to land management practices (Yamakoshi & Leblan 2013; Leblan & Soiret 2021), and some may be older than the usual 20–25-year-old oil palms at large-scale plantations (Jourdan & Rey 1997). As this is a resource shared by the people and chimpanzees of Bossou (Humle & Matsuzawa 2004; Hockings et al. 2009), there is the risk of anthropic input of phytolith assemblages produced. This will have to be carefully factored in to ensure initial interpretations of phytolith assemblages from directed input can be done that are independent from anthropic input.

In terms of analyses of production of phytolith assemblages at different sediment depths in relation to the nut-cracking sites at Bossou, AIF/g of the various depths appears to indicate some possible variation in phytolith productivity, with non-nut-cracking sites having a more similar AIF/g of spheroid echinates across the depths sampled, compared to the dormant and active nut-cracking sites which visually appear to be more productive in near surface sediment depths (Fig. 6). We could not determine this statistically in this study due to sample size, but again, an increased sample size and study of spheroid echinates across the different depths would clarify if variation of phytolith productivity across the depths results from nut-cracking activity by chimpanzees. The movement of phytoliths from the surface down to deeper sediments has understandable received long-standing attention and has indicated how burning, size of phytoliths, pH, bioturbation, water exposure and draining, and soil particle size all support Hart & Humphreys (1997) mobile phytolith hypothesis (Rovner 1986; Fishkis et al. 2009, 2010; Cabanes et al. 2011) and whether we observe gradual decrease or increase in phytolith movement (Hart & Humphreys 2003). Does the physical impact of repeated use of percussive technology, cause smaller spheroid echinate phytoliths to move further down in deeper sediments? Although work has been done to explore the surfaces of percussive technology such as grinding stones and the use of large pestle and mortars for recent humans (Hayes et al. 2018; Tsartsidou & Ktsakis 2020), as well as for stone tools of our ancestors (Domínguez-Rodrigo et al. 2001), no published literature could be found to address this question.

Our preliminary data on approximated 2D area of the spheroid echinates as well as perimeter and maximum diameter all indicated intra-specific differences in the size of this phytolith morphotype in the nut and leaflet of the oil palm. This is rather a crude approximation compared to the detailed work done using phytolith morphometrics (Ball et al. 2016; Yansheng et al. 2019) and on globular and palm phytoliths (Fenwick et al. 2011; Benvenuto et al. 2015; Huisman et al. 2018; Witteveen et al. 2022), however, published data on detailed morphometrics of oil palm phytoliths does not seem available. Should there be intra-specific differences between nut endocarp phytoliths and that of other parts of the oil palm in that the former are generally larger as indicated in this study, this would have major implications for interpretations of phytolith assemblages in sediments associated with processing oil palm nuts by both human and non-human taxa. Our next step is to measure a proportion of the spheroid echinates in existing sediment samples, particularly comparing phytolith 2D area and maximum diameter for surface sediments at the nut-cracking and non-nut-cracking sites; in addition, to collect additional samples as discussed and to also carry out further measurements across multiple parts of the oil palms of Bossou Forest.

There are multiple future directions in which to take this work, some described already in relation to increasing sample size at Bossou Forest, collecting samples at additional sites where chimpanzee nut-cracking has occurred, completing an in-depth study of phytolith assemblages at different sediment depths at Bossou, particularly in sediments in localities where repeated use of percussive technology takes place, and obtaining detailed measurements on oil palm phytolith morphometrics. There are additional future directions to explore for the directed input of oil palms by nut-cracking activity of the Bossou chimpanzees, such as how the use of percussive technology impacts the structure of phytoliths that are being processed (i.e. if there would be an increase in partial spheroid echinates pre- and post-deposition due to possible damage from the crushing the oil palm nut). The scope to explore directed input from other aspects of their complex foraging strategies such as the insectivory component of their diet (using perishable tools for ant-dipping and termite fishing) also awaits further attention. Finally, there are *Raphia* palms in Bossou Forest where the chimpanzees feed on the gum as a result of visiting farms and causing crop damage (Hockings et al. 2009). There phytoliths could also be studied to compare with oil palms phytoliths.

## Conclusion

In the study of directed input by non-human taxa and the interpretation of phytolith assemblages produced, this work has immense potential to explore ecological spaces shared by human and non-human taxa, such as primates and highlight the mutual roles they play (Fuentes 2012) and highlight the unique cultural heritage of our close relatives to work towards ensuring the survival of as many primates as possible (Estrada et al. 2017; Kühl et al. 2019; Hockings and McLennon 2019; Carvalho et al. 2022). This is ever more crucial and urgent for the Bossou chimpanzees who remain biogeographically isolated from most neighbouring chimpanzee communities. Furthermore, it also has potential to contribute and build upon work done at existing primate archaeological sites for chimpanzees such as that in Tai Forest, Ivory Coast (Mercader et al. 2000; 2007), but also provide insight into the interpretation of phytolith assemblages at nut-cracking sites yet to be discovered for extirpated chimpanzees of West Africa (Smith et al. 2010) and at other sites where such directed plant input is observed for primates, for example for the bearded capuchins at Boa Vista, Brazil who use stone hammers and wooden and boulder anvils to crack open various palm nuts, or the long-tailed macaques who crack open oil palm nuts in Ao Phang-Nga National Park, Thailand (Luncz et al. 2017). Finally, it highlights that there are many future directions needed to expand work on interpreting phytolith assemblages from directed input of non-human taxa.

## Acknowledgements

We thanks Fromo Dore and Pascal Goumi for support in the field and the Direction Nationale de la Recherche Scientifque, the Institut de Recherche Environmentale de Bossou (Guinea), and the Kyoto University Primate Research Institute, for research permissions. CP was funded by the NRF DST Centre of Excellence in Palaeosciences and the Leverhulme Trust. CP is the main author and was responsible for conceptualisation, methodology, and writing of most of the manuscript. KAW was responsible for most of the primary data collection, and logistical support in the field. MB was responsible for logistical support for laboratory work in the Palynology and Phytolith Lab at the Evolutionary Studies Institute, University of the Witwatersrand and supervision of CP during this postdoctoral work. We thank Katherine Willis for allowing us to analyse sediment samples in the Oxford Long-Term Ecology Lab, in Dept. of Biology, University of Oxford as well as Steve Boreham and Chris Rolfe for allowing us to analyse sediment samples in the Geography Science Laboratories, University of Cambridge.

## Notes

### Competing Interest Statement

The authors have declared no competing interest.

