## Supplemental Material 1 for "Interpreting phytoliths assemblages at chimpanzee (*Pan troglodytes verus*) nut-cracking sites in Bossou Forest, Guinea": SM1.pdf

| Site a | Munsell | pH | Particle size |
|---|---|---|---|
| 0-2 cm | 7.5YR 2.5/3 | 5.2 | 30.56 |
| 2-5 cm | 7.5YR 3/4 | 5.7 | 14.12 |
| 5-10 cm | 7.5YR 3/5 | 5.1 | 256.93 |
| Site b | | | |
| 0-2 cm | 5YR 2.5/2 | 5.2 | 39.21 |
| 2-5 cm | 5YR 3/4 | 5.0 | 153.76 |
| 5-10 cm | 7.5YR 2.5/3 | 5.3 | 210.41 |
| 10-15 cm | 5YR 3/4 | 5.7 | 43.87 |
| Remainder | 2.5YR 3/6 | 5.6 | 284.17 |
| Site c | | | |
| 0-2 cm | 5YR 3/4 | 5.1 | 155.31 |
| 2-5 cm | 5YR 3/3 | 5.5 | 31.09 |
| 5-10 cm | 5YR 3/4 | | 17.51 |
| 10-15 cm | 5YR 3/4 | 5.6 | 12.42 |
| Site d | | | |
| 0-2 cm | 7.5YR 2.5/3 | 5.0 | 67.03 |
| 2-5 cm | 7.5YR 2.5/3 | 5.0 | 10.92 |
| 5-10 cm | 5YR 3/3 | 5.5 | 25.28 |
| Remainder | 5YR 3/4 | 5.2 | 20.7 |
| Site e | | | |
| 0-2 cm | 5YR 2.5/2 | 5.3 | 22.81 |
| 2-5 cm | 7.5YR 2.5/3 | 5.4 | 15.84 |
| 5-10 cm | 5YR 3/3 | 5.3 | 60.89 |
| 10-15 cm | 5YR 3/4 | 5.6 | 48.59 |
| Remainder | 5YR 3/4 | 5.6 | 52.36 |
| Site f | | | |
| 0-2 cm | 5YR 3/4 | 5.7 | 26.66 |
| 2-5 cm | 5YR 3/4 | 5.3 | 9.03 |
| 5-10 cm | 5YR 3/4 | 4.9 | 11.19 |
| 10-15 cm | 5YR 3/4 | 5.2 | 44.26 |

| 0-2cm | | pH | 2-5cm | | pH | 5-10cm | | pH | 10-15cm | | pH |
|---|---|---|---|---|---|---|---|---|---|---|---|
| 30.56 | silt | 5.2 | 14.12 | silt | 5.7 | 256.93 | sand | 5.1 | | | |
| 39.21 | silt | 5.2 | 153.76 | sand | 5 | 210.41 | sand | 5.3 | 43.87 | silt | 5.7 |
| 155.31 | sand | 5.1 | 31.09 | silt | 5.5 | 17.51 | silt | | 12.42 | silt | 5.6 |
| 67.03 | sand | 5 | 10.92 | silt | 5 | 25.28 | silt | 5.5 | | | |
| 22.81 | silt | 5.3 | 15.84 | silt | 5.4 | 60.89 | silt | 5.3 | 48.59 | silt | 5.6 |
| 26.66 | silt | 5.7 | 9.03 | silt | 5.3 | 11.19 | silt | 4.9 | 44.26 | silt | 5.2 |

sand 2000-63
silt 63-2
clay <2

pH range 4.9 to 5.7 (moderately acidic)
