## Supplemental material 2 for "Interpreting phytoliths assemblages at chimpanzee (*Pan troglodytes verus*) nut-cracking sites in Bossou Forest, Guinea": SM2.pdf

| Collected 2018 | | Slide Transect 1 | Slide Transect 2 | Slide Transect 3 | Slide Transect 4 | Slide Transect 5 | Slide Transect 6 | Slide Transect 7 | Slide Transect 8 | Slide 9 |
|---|---|---|---|---|---|---|---|---|---|---|
| Sediment sample | | | | | | | | | | |
| T1 0-5 | DNC | 16 | 27 | 13 | 19 | 20 | 5 | 16 | 23 | |
| T1 10-15 | DNC | 33 | 31 | 18 | 4 | 15 | 8 | 11 | 9 | |
| T2 0-5 | DNC | 19 | 13 | 20 | 9 | 18 | 2 | 9 | 6 | |
| T2 5-10 | DNC | 27 | 39 | 14 | 13 | 9 | 46 | 28 | 35 | |
| T2 10-15 | DNC | 16 | 23 | 14 | 26 | 13 | 15 | 22 | 16 | |
| T3 0-5 | NNC | 12 | 9 | 11 | 5 | 18 | 16 | 6 | 11 | |
| T3 5-10 | NNC | 9 | 6 | 7 | 15 | 10 | 11 | 7 | 9 | |
| T3 10-15 | NNC | 13 | 3 | 13 | 5 | 4 | 5 | 3 | 7 | |
| T4 0-5 | NNC | 7 | 7 | 10 | 6 | 13 | 6 | 10 | 9 | |
| T4 5-10 | NNC | 15 | 26 | 28 | 19 | 21 | 26 | 22 | 28 | |
| T4 10-15 | NNC | 5 | 3 | 3 | 3 | 1 | 4 | 3 | 5 | |
| T5 0-5 | ANC | 38 | 17 | 11 | 18 | 21 | 28 | 19 | 16 | |
| T5 5-10 | ANC | 9 | 19 | 20 | 36 | 20 | 19 | 19 | 13 | |
| T5 10-15 | ANC | 27 | 12 | 27 | 14 | 26 | 10 | 16 | 13 | |
| T6 0-5 | ANC | 6 | 2 | 8 | 13 | 6 | 6 | 7 | 8 | |
| T6 5-10 | ANC | 15 | 16 | 24 | 23 | 9 | 14 | 9 | 15 | |
| T6 10-15 | ANC | 17 | 9 | 16 | 20 | 13 | 9 | 13 | 3 | |

Plant — Transect slide number different as different sliides used during microscopy

| Plant | Slide Transect 1 | Slide Transect 2 | Slide Transect 3 | Slide Transect 4 | Slide Transect 5 | Slide Transect 6 | Slide Transect 7 | Transect 8 |
|---|---|---|---|---|---|---|---|---|
| LEAF 1 | 118 | 77 | 118 | 63 | 70 | 202 | 131 | 135 |
| LEAF 2 | 162 | 113 | | | | | | |
| NUT 1 | 11 | 23 | 18 | 22 | 24 | 32 | 20 | 19 |
| NUT 2 | | | | | | | | |
