## Supplemental material 3 for "Interpreting phytoliths assemblages at chimpanzee (*Pan troglodytes verus*) nut-cracking sites in Bossou Forest, Guinea": SM3.pdf

| | Nut 2D area sq µm | Leaflet 2D area sq µm | Nut perimeter µm | Leaflet perimeter µm | Nut length µm | Leaflet length µm |
|---|---|---|---|---|---|---|
| 1 | 108.6 | 30.4 | 42.3 | 23.1 | 13.2 | 7 |
| 2 | 28.3 | 35.1 | 28.4 | 24.6 | 5.4 | 6.8 |
| 3 | 189.35 | 79.2 | 53.4 | 35.6 | 16.8 | 11.6 |
| 4 | 87.9 | 26 | 41.1 | 21.2 | 11.4 | 7.2 |
| 5 | 18 | 20.5 | 20.7 | 20.2 | 5 | 6.4 |
| 6 | 55.1 | 34.9 | 31.7 | 25 | 8.2 | 7.4 |
| 7 | 128.1 | 21.3 | 45.3 | 19.8 | 14.2 | 6.2 |
| 8 | 114.6 | 20.7 | 44.1 | 20.2 | 13.8 | 6.6 |
| 9 | 110.9 | 18.3 | 41.6 | 18.4 | 12.8 | 6.3 |
| 10 | 36 | 33.5 | 25.4 | 23.9 | 8.3 | 8.2 |
| 11 | 137.4 | 40.4 | 51.5 | 26.6 | 14.7 | 8.5 |
| 12 | 172 | 61.2 | 60.9 | 31.7 | 16.8 | 9.5 |
| 13 | 95.9 | 17.9 | 42.1 | 18 | 11.4 | 4.9 |
| 14 | 143.9 | 40 | 50.8 | 26.6 | 16 | 8.4 |
| 15 | 135.8 | 40.3 | 49.6 | 26.3 | 15.7 | 9 |
| 16 | 381.4 | 24.4 | 78.3 | 22.7 | 23.7 | 8.4 |
| 17 | 177.55 | 27.6 | 54.4 | 23.2 | 16.9 | 7.4 |
| 18 | 189.4 | 25.4 | 60.3 | 21.1 | 17.2 | 6.4 |
| 19 | 128.8 | 34.2 | 49.1 | 24.9 | 14.2 | 7.9 |
| 20 | 163 | 27.1 | 57 | 21.4 | 14.7 | 6.4 |
| 21 | 186.6 | 21.7 | 60.1 | 19.5 | 16.1 | 6.3 |
| 22 | 404.6 | 44.5 | 80.7 | 30.2 | 25.1 | 9.5 |
| 23 | 175.7 | 60.8 | 59.2 | 31.9 | 16.6 | 9.4 |
| 24 | 208.8 | 46.55 | 60.7 | 29.7 | 18.4 | 9.2 |
| 25 | 217 | 25.3 | 64.1 | 21.3 | 18.3 | 7.3 |
| 26 | 208.5 | 30.7 | 58.5 | 22.8 | 17.7 | 7.2 |
| 27 | 103.2 | 83.7 | 48.8 | 38.7 | 12.6 | 10.9 |
| 28 | 206.6 | 33.75 | 60.5 | 26.4 | 17.3 | 8.3 |
| 29 | 149.2 | 26.8 | 54.6 | 21.3 | 14.6 | 6.8 |
| 30 | 288.2 | 36.6 | 69.3 | 26.4 | 19.9 | 8.1 |
